# Structural determinants of endopilus assembly, stability and functional specificity in bacterial type II secretion

**DOI:** 10.64898/2026.04.10.717868

**Authors:** Maylis Lejeune, Stefaniia Ivashchenko, Régine Dazzoni, Benjamin Bardiaux, Ravi R. Sonani, Matthijn Vos, Theis Jacobsen, Edward H. Egelman, Michael Nilges, Olivera Francetic, Vladimir E. Shevchik, Nadia Izadi-Pruneyre

## Abstract

Gram-negative bacteria employ the type II secretion system (T2SS) to transport folded protein effectors across the outer membrane. This multiprotein nanomachine assembles the endopilus, a periplasmic helical polymer composed of one major and four minor pilin subunits. Endopili are structurally related to type IV pili but exhibit distinct features including a conserved calcium-binding site stabilizing their major pilin subunits. Endopilus polymerisation is coupled to substrate translocation through a dedicated outer membrane channel.

To investigate the structural basis of endopilus assembly, stability and functional specificity, we performed a comparative analysis of the Out T2SS from the plant pathogen *Dickeya dadantii* and the Pul T2SS from the human pathogen *Klebsiella oxytoca.* Although their major pilins OutG and PulG share over 77% sequence identity, these bacteria differ markedly in ecological niche and range of secreted effectors.

We report here the NMR structure of calcium-bound OutG monomer and cryo-EM structures of OutG and PulG endopili at 3.6 Å resolution. The integration of structural, mutational, and biophysical analyses with *in vivo* assays, allowed us to identify the molecular determinants of secretion specificity and endopilus stability. These findings demonstrate how subtle sequence variations in conserved nanomachines have evolved to optimize their function and adapt to their local environments.

## Introduction

In Gram-negative bacteria, type II secretion systems (T2SS) mediate the transport of specific folded proteins across the outer membrane^1^. Secreted substrates or effectors include lytic enzymes, toxins, adhesins or cytochromes with key roles in virulence, competition and niche adaptation^2^. T2SSs are multicomponent trans-envelope nanomachines, that polymerize periplasmic fibres known as endopili (previously called pseudopili) essential for protein secretion (**Figure 1A**). The endopilus core is a helical polymer of major pilins, generically named GspG, and four low abundance (minor) pilins GspH, GspI, GspJ and GspK, which initiate fibre assembly and are predicted to cap its distal end^3–5^. Endopilus assembly is coupled to the secretion of effector proteins to the extracellular milieu across the dedicated outer membrane channel, the secretin^6^; however, the mechanism of this coupling remains unknown. T2SSs are members of the type IV filament superfamily of prokaryotic nanomachines^7^, which also includes type IV pili (T4P), long surface filaments that promote bacterial adhesion, surface motility and macromolecular transport^8,9^. Consistent with their evolutionary similarity to T4P, overexpression of T2SS genes causes artificial extension of endopili into long filaments displayed on the bacterial cell surface^10^ (**Figure 1A**). This feature has facilitated the biochemical, structural and functional analyses of these extended filaments, commonly referred to as T2SS pili. Combined modelling and biochemical studies have further shown that T2SS pili share a similar architecture and helical subunit organization with T4P, notably with the T4aP class^11,12^. However, unlike the major T4a pilins, which are typically constrained by one or two disulfide bonds, the T2SS major pilins are stabilized by a calcium ion bound to a conserved region close to their C-terminus – the calcium binding site^13^.

**Figure 1:**
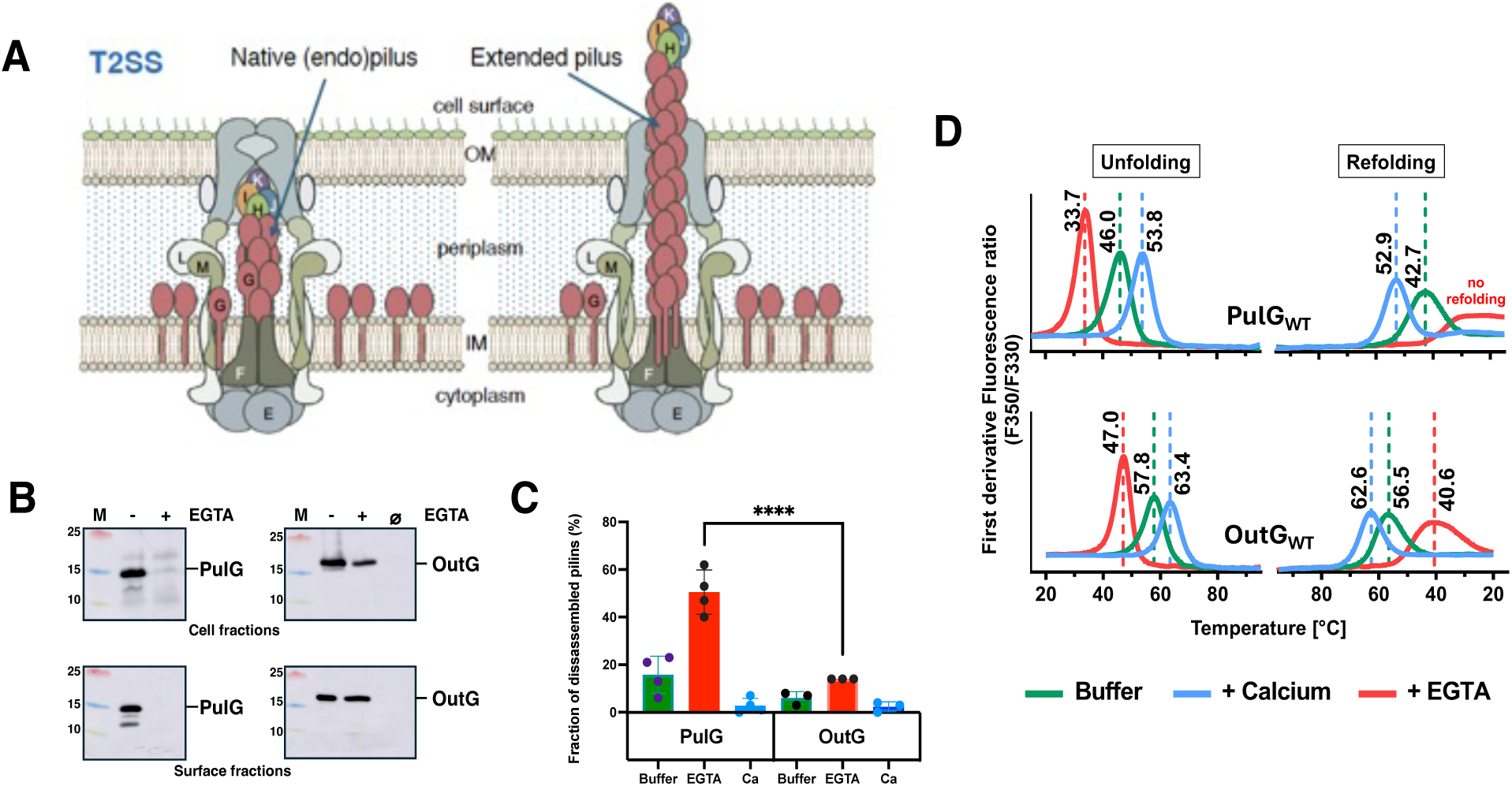
Role of calcium in the stability of OutG and PulG subunits and pili. **A.** Schematic representation of T2SS organisation showing endopilus (left) and extended pilus (right). The major pilin GspG is shown in brown. **B.** Immunodetection of PulG and OutG in the cell and surface fractions of *E. coli* harbouring *K. oxytoca* T2SS cultured on LB plates in the absence or presence of EGTA. **C.** Pilus stability *in vitro* assay. Isolated pili were incubated 1 hour at 60°C in 50 mM HEPES, 50 mM NaCl buffer, and buffer supplemented with 5 mM EGTA or 5 mM CaCl_2_. Pili and dissociated pilin subunits were separated by ultracentrifugation and quantified after SDS-PAGE and Coomassie blue staining (Figure S1). Bar heights show mean levels of disassembled pilus fractions measured from 3 and 4 independent experiments for OutG and PulG, respectively. Error bars indicate standard deviation. **** (p<0.001). ***D*.** Unfolding (left) and refolding (right) curves of purified PulG and OutG periplasmic domains measured by NanoDSF in buffer (green curves), with added CaCl_2_ (blue) or EGTA (red). The first derivative of 350nm/330nm fluorescence with respect to temperature is represented. The transition temperature (Tm) values are indicated above each curve.

The first experimentally determined structure of a T2SS pilus, obtained by integrating the nuclear magnetic resonance (NMR) structure of the major pilin PulG from *Klebsiella oxytoca* with the 5 Å resolution cryo-electron microscopy (cryo-EM) map of the pilus^14^, revealed the organization of calcium binding sites at the pilus surface. However, to improve the cryo-EM map resolution, a non-native PulG variant stabilized by an intramolecular disulfide bond^14^ was used. *In vitro* and *in vivo* analyses further confirmed the key role of calcium in pilin monomer subunit folding and in the stability of the assembled PulG pilus^14^.

The T2SS endopilus promotes the secretion of specific effectors, and may play a role in their selection and recruitment through direct interaction^1^. To investigate how endopilus structure governs functional specificity, here we performed a comparative structural and functional analysis of the T2SS pili from *K. oxytoca* (a human pathogen) and *Dickeya dadantii* (a plant pathogen). These systems differ in their host habitat and range of effectors: *K. oxytoca* Pul system secretes only a starch-degrading enzyme pullulanase^15^, whereas *D. dadantii* Out T2SS secretes ∼20 plant cell wall-degrading enzymes (primarily polysaccharide lyases) to drive virulence^16^. Despite this marked functional difference, their major pilins PulG and OutG share over 77% sequence identity, making them an ideal model system to dissect the molecular basis of major pilin and pilus specificity.

Towards this goal, we determined the structures of the calcium-bound OutG monomer and pilus by solution NMR and cryo-EM, respectively. To compare these structures with those of *K. oxytoca* PulG, we also solved the cryo-EM structure of the wild type *K. oxytoca* PulG pilus. The two cryo-EM maps of PulG and OutG pili at 3.6 Å resolution revealed new features including the fine details of the calcium-binding site. By integrating structural, mutational and biophysical analyses with *in vivo* phenotypic assays, we identified the distinct sequence and structural determinants of major pilins that govern T2SS secretion specificity and endopilus stability.

## Results

### Calcium dependence of PulG and OutG stability and pilus assembly

*Escherichia coli* carrying a complete set of genes that encode the *K. oxytoca* T2SS assembles PulG pili on the cell surface when grown on agar plates (**Figure 1B**, left). Supplementing the growth medium with the calcium chelator ethylene glycol-bis (β-aminoethyl ether)-*N*,*N*,*N*’,*N*’-tetraacetic acid (EGTA) leads to marked reduction in cellular levels of PulG and abolishes piliation, as shown previously^14^. Replacing *pulG* with *outG* encoding the major pilin of *D. dadantii* OutG in the same strain led to efficient formation of OutG pili on the bacterial surface. In contrast to PulG, the cellular levels of OutG were marginally affected and the bacteria assembled comparable amounts of OutG pili regardless of EGTA presence (**Figure 1B**, right).

We also compared the stability of OutG and PulG pili isolated from these strains by quantifying their disassembly after incubation at 60°C in buffer, or buffer containing 5 mM CaCl_2_ or 5 mM EGTA (**Figure 1C, S1)**. OutG pili exhibited remarkable stability, showing no detectable disassembly in buffer or buffer supplemented with CaCl₂, and only minimal dissociation (< 10%) upon EGTA treatment. In contrast, PulG pili displayed moderate instability in buffer (∼15%) and significantly greater disassembly in EGTA, with more than 50% of pilins dissociated from the filaments (**Figure 1C**).

To test whether the difference in stability between PulG and OutG pili is governed by the intrinsic stability of their subunits, we assessed the behaviour of isolated pilins *in vitro*. To this end, we purified soluble periplasmic domains of PulG and OutG lacking the hydrophobic transmembrane segments and investigated their stability by Nano-Differential Scanning Fluorimetry (NanoDSF). This method monitors protein unfolding, refolding and conformational transitions across a temperature gradient by tracking the intrinsic fluorescence of tryptophan residues. PulG and OutG at the same concentration were analysed under three conditions: i) native, as extracted and purified from bacteria; ii) supplemented with CaCl_2_ or iii) with EGTA. Native OutG was more stable than PulG (Tm 63.4°C versus 53.8°C respectively). The stability of both proteins was similarly affected by depletion of calcium ions with EGTA, with a decrease in Tm of 15-20°C. However, in the absence of calcium, PulG was unable to refold, demonstrating a strict calcium dependence, while OutG refolded at 40.6°C, albeit partially, as indicated by a broad peak (**Figure 1D**). Thus, the *in vitro* behaviour of PulG and OutG correlates with that observed *in vivo,* suggesting that it is caused by intrinsic properties of the two proteins.

### Structure of OutG pilin and comparison with PulG

In a previous study, we showed that the native PulG monomer, produced in the periplasm, was in a calcium-bound state^14^. Similarly, the NMR spectral signature of the OutG monomer indicates a calcium-bound state, as its ^1^H-^15^N HSQC spectrum reflecting the overall protein fold, is identical to that of OutG in the presence of added calcium. Moreover, calcium addition to the EGTA-treated OutG sample restored the NMR spectral signature of native OutG (**Figure S2**). Both calcium and EGTA induced local spectral modifications (**Figure 2A**). We analysed the chemical shift perturbation (CSP) of backbone amide resonances between the calcium-bound and calcium-free states (**Figure 2A, 2B**). Residues 114-126 show the largest spectral perturbations (CSP > 0.2 ppm) upon calcium binding, indicating that their chemical environment was modified either by direct coordination *via* their backbone or their side chain, or by local conformational changes. Residues D92-L100 located just upstream of the calcium binding site were also affected, but with lower CSP (< 0.1 ppm) (**Figure 2B, 2D, 2E**).

**Figure 2.**
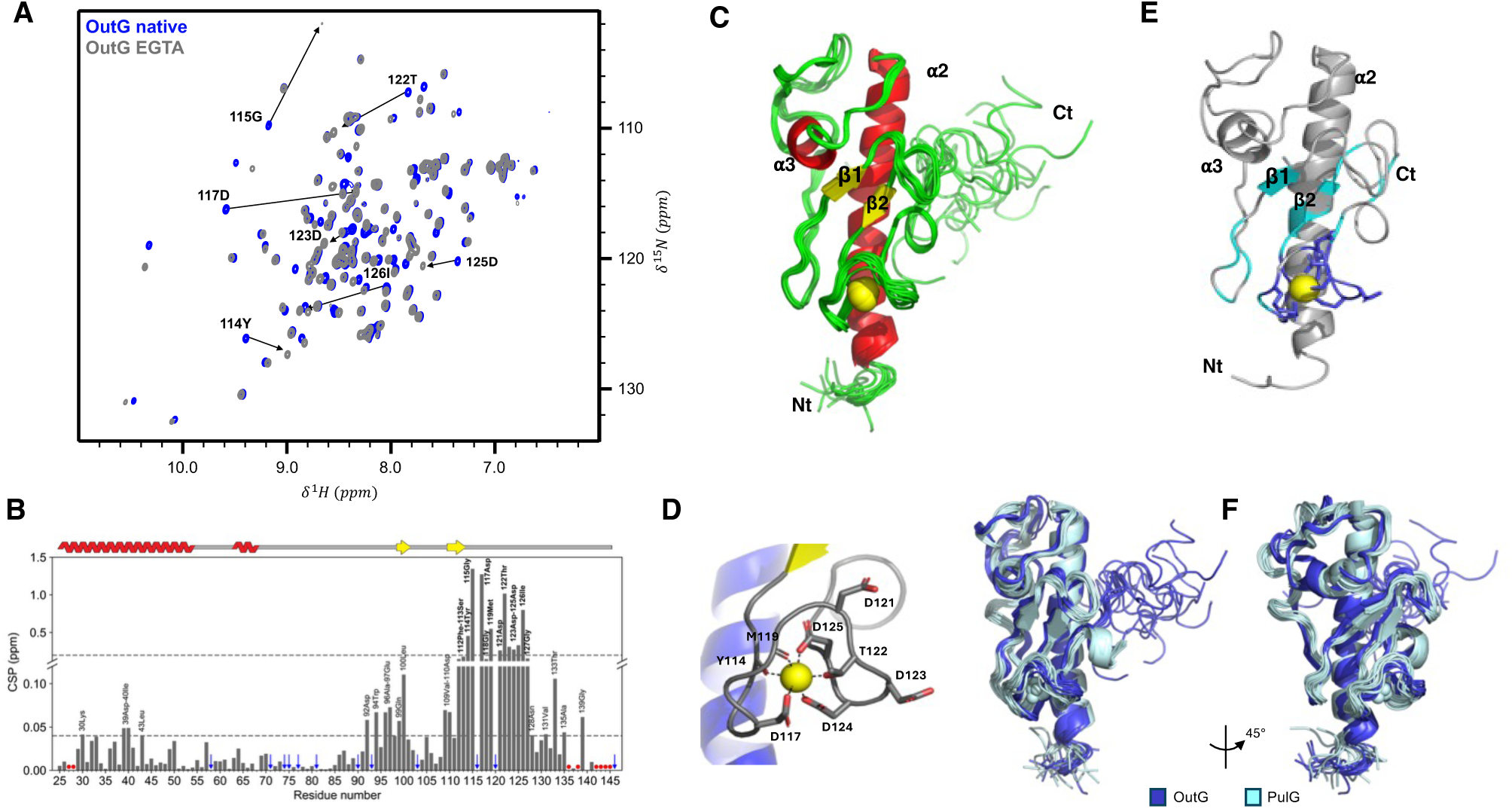
Structure of the OutG pilin in the calcium-bound state and comparison with PulG. **A**. Identification of OutG calcium binding site. Superimposed ^1^H-^15^N HSQC NMR spectra of native OutG pilin at 0.1 mM, purified in 50 mM HEPES,100 mM NaCl, pH 7 buffer (blue), and in buffer supplemented with 10 mM EGTA (grey). Resonances corresponding to calcium binding residues in both forms are labelled. **B**. Chemical shift perturbation (CSP) upon calcium binding (red dots = unassigned, blue arrow = proline not visible in ^1^H-^15^N HSQC). Residues with CSP > 0.04 ppm are labelled (in bold for the calcium binding site). Secondary structures are shown on top. **C**. OutG solution NMR structure ensemble coloured by secondary structure elements. Calcium is shown as a yellow sphere. **D.** Coordination of Ca^2+^ in the OutG NMR ensemble. **E.** OutG structure coloured by CSP as in B (cyan > 0.04 ppm, blue > 0.2 ppm); **F**. Superimposition of OutG (blue) and PulG (PDB 5O2Y, cyan) solution structure ensembles (Cα RMSD 2 Å).

We determined the solution structure of the periplasmic domain of OutG by NMR spectroscopy **(Figure 2C)**. For structure calculation, 1593 distance restraints were derived from 3D ^15^N- and ^13^C-NOESY experiments using ARIAweb^17^, together with 173 backbone dihedral angle restraints predicted from secondary chemical shits^18^. Structure and restraints statistics for the OutG NMR ensemble are given in **Table S1**.

The OutG monomer adopts a lollipop-like architecture characterized by an α/β fold including a long N-terminal helix (K28-D53) followed by a short helix (G63-V68) and a two-stranded antiparallel β-sheet (Q99-V101; D110-F112) (**Figure 2C**). The C-terminal extension (V131-P146) is disordered^19^. This structure is highly similar to that of PulG monomer in solution (PDB 5O2Y)^14^, with a Cα RMSD of 2.2 Å calculated over residues M25 to G132 (**Figure 2F**). Consistent with CSP data, calcium is coordinated by oxygen atoms of the side chains of D117, T122 and D125 as well as backbone carbonyl oxygens of Y114 and M119. Interestingly, in the OutG NMR ensemble, 2 out of 10 conformers show coordination of calcium also by D124 side chain. Side chains of D121 and D123 point away from the calcium ion and are not part of the coordination **(Figure 2D)**. Overall, the conformation of the calcium binding site in OutG is similar to that of PulG^14^.

### Structure of OutG and PulG pili

To investigate how the interactions between pilin subunits contribute to the remarkable stability of assembled OutG pilus, we determined its structure by cryo-EM. The OutG pili sample was prepared as described in Materials and Methods. In cryo-EM micrographs, OutG pili appeared as ∼8 nm-wide filaments displaying a sinuous morphology (**Figure 3A**). The final 3D reconstruction at ∼3.6 Å resolution allowed us to build an atomic model of the OutG pilus in the calcium-bound state (**Figure 3C, 3E**). The helical parameters of the 1-start filament (axial rise 10.8 Å and twist angle 84.1°) are similar to the previously obtained PulG pilus structure^14^ (**Table S2**). Since the previous PulG pilus structure was determined for a variant stabilized with a disulfide bridge (H106C-W129C) and at a lower resolution (∼5 Å), we also determined the structure of the wild-type PulG pilus by cryo-EM at a resolution of ∼3.6 Å (**Figure 3B, 3D**). The wild-type PulG pilus adopts a slightly more compact form with a smaller axial rise (10.5 Å) compared to the OutG pilus. As observed in the previous PulG model and in T4P structures^20,21^, the stem region of the pilin presents a discontinuous N-terminal helix, with a non-helical, melted region between S18 and G26 (**Figure 3E, 3F**). The C-terminal extension of OutG (T133-P146) was invisible in the cryo-EM reconstruction and was thus not possible to model, similar to PulG where the last 4 residues (I131-K134) are not resolved, certainly due to a conformational flexibility. Multiple conformations of this region were observed in NMR spectra of the monomer subunit^19^. OutG and PulG monomeric pilin structures within the pilus are virtually identical with a Cα RMSD of 0.9 Å (130 residues aligned, **Figure 3F**).

**Figure 3:**
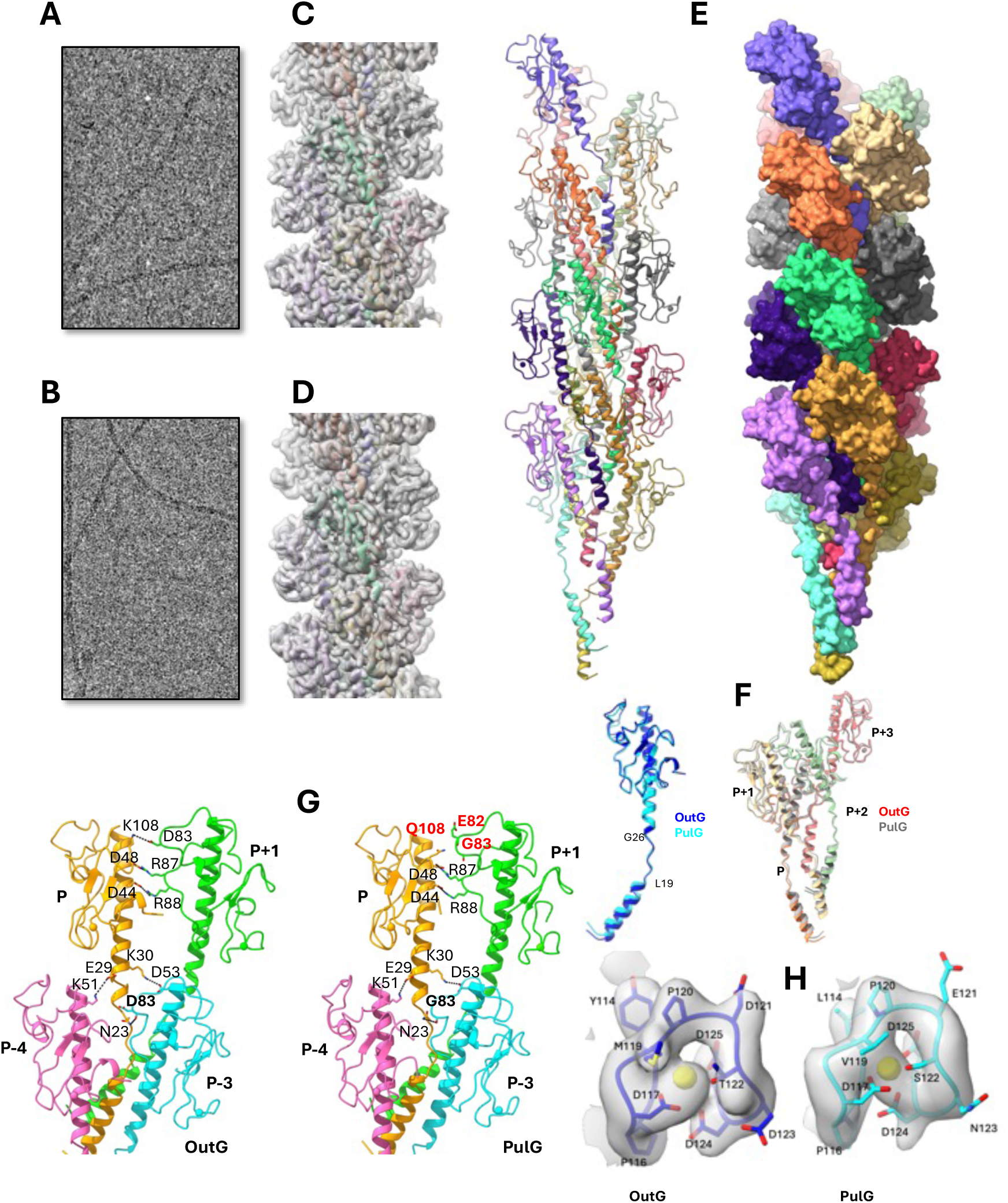
OutG and PulG pilus structures. **(A, B)** OutG and PulG pili cryo-EM micrographs **(C, D)** OutG and PulG cryo-EM reconstructions**. E.** OutG pilus structure represented as cartoon (left) and surface (right), coloured by subunit**. F.** Superimposition of OutG (blue) and PulG (cyan) monomers (left) and 4-subunit oligomers (right). **G.** Inter-subunit contacts in OutG (left) and PulG (right) pilus structures. Subunits P, P+1, P-3 and P-4 along the 1-start helix are shown. **H.** EM densities of calcium binding sites in OutG (left) and PulG (right) pili. Calcium is shown in yellow.

The overall topology of OutG and PulG pili is highly similar (Cα RMSD 1.1 Å over four consecutive subunits). In both structures, each subunit interacts with three subunits above, P+1, P+3 and P+4, along the helical filament, and three below in a symmetrical way (**Figure 3F, 3G**).

The two salt bridges involving the negatively charged residues E44 and D48 of subunit P and arginine residues 87 and 88 in subunit P+1 are found in both pili. These four residues are highly conserved in T2SS pilins, and their interactions have been shown to be essential for pilus assembly and protein secretion^11^. In OutG pilus, this interface is reinforced by a third salt bridge between D83 from subunit P with K108 from subunit P+1. In PulG, the equivalent residues G83 and Q108 are not involved in any interaction, and the side chain of neighbouring E82 does not contribute to this interface (**Figure 3G**).

Stabilizing interactions between residues K51 and E29 from subunits P and P+4 respectively, are present in both OutG and PulG pili. In both pilus structures, D53 is close to residue K30, suggesting a stabilizing contact at the interface between P and P-3 subunits. These contacts maintain pilus stability but are not essential for protein secretion, as demonstrated previously^11,12^. Notably, the increased resolution of the PulG wild type pilus cryo-EM reconstruction revealed a contact between the conserved N23 side chain in the melted portion of the N-terminal helix and the backbone carbonyl oxygen of residue 83 (D83 in OutG, G83 in PulG). This interaction, at the interface of subunit P with P+3 and P-3 subunits, also present on the OutG pilus, is most likely formed upon pilus assembly and may contribute to the stabilization of this extended unfolded region of the filament (**Figure 3G**).

Interestingly, all sequence differences between OutG and PulG map onto the accessible surface of the periplasmic domain of the pilin subunits when assembled in pili, except for Q82, D83, K108 in OutG and E82, G83 and Q108 in PulG (**Figure 4A, 4B)**. This residue variation results in a more negative surface potential for the PulG pilus compared to OutG (**Figure 4C**).

**Figure 4.**
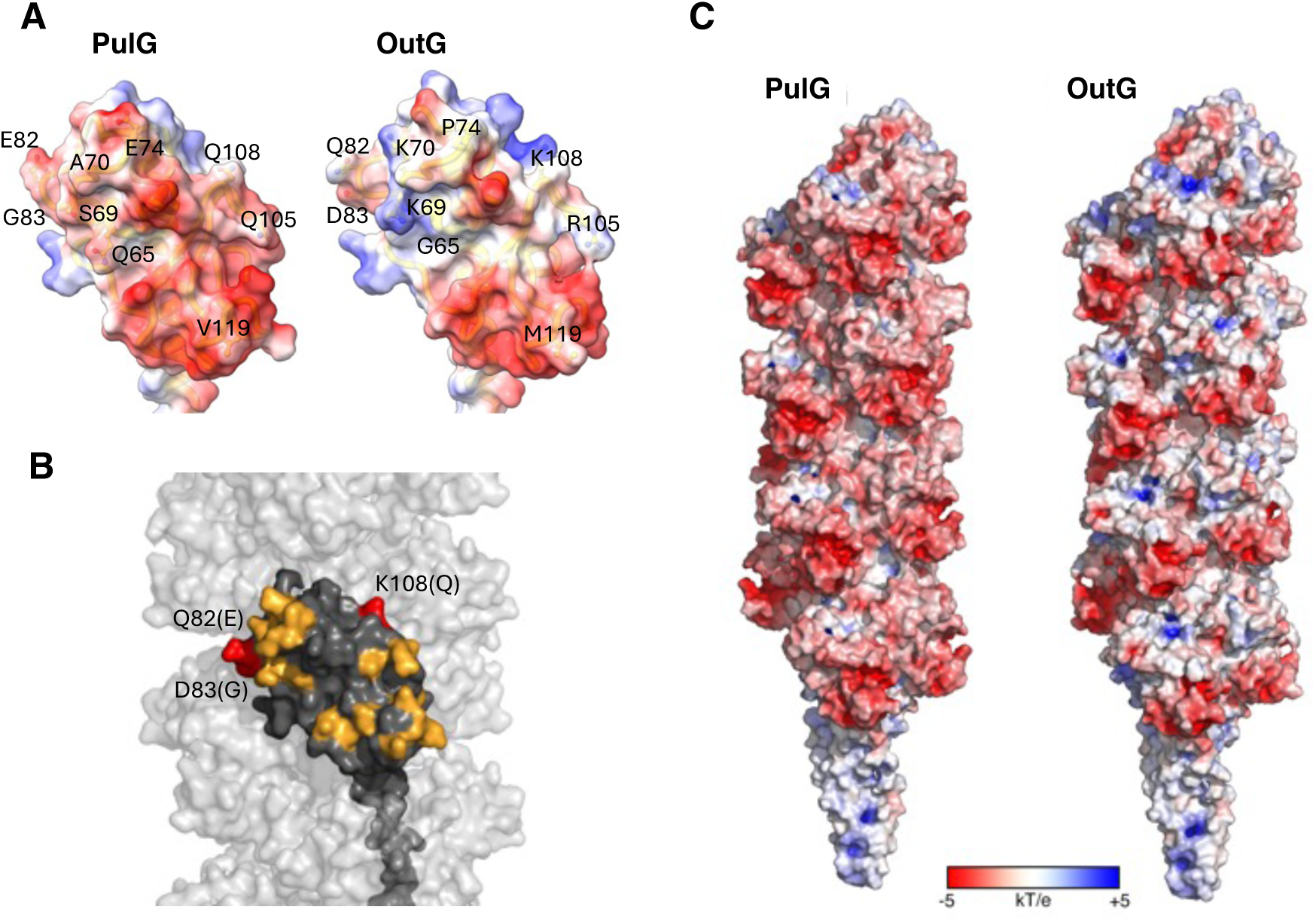
OutG and PulG surface differences. **A.** Surface electrostatic potential of PulG and OutG monomer subunits (periplasmic domain). Residues that differ between the two proteins are labelled. **B.** Surface representation of an OutG subunit (in dark grey) within the pilus (light grey). Variable residues are highlighted in orange (all surface-exposed), except Q82(E), D83(G), and K108(Q), which are buried and marked in red. The corresponding residues in PulG are indicated in parentheses. **C.** Surface electrostatic potential of PulG and OutG pili structures.

In cryo-EM reconstructions, density for calcium ions is observed in both pili at the calcium-binding site, which is exposed at the pilus surface **(Figure 3H).** Superposition of this loop reveals a highly similar organization of residues coordinating the calcium ion (**Figure S3**). As observed by NMR, the side chains of the conserved residues D117 and D125 (and potentially D124) are the main calcium coordinating ligands, along with Y114, M119 and T122 in OutG (L114, V119 and S122 in PulG).

### Sequence and structure determinants of calcium-dependent pilin stability

Comparative structural analyses of OutG and PulG monomer subunits and their assembled pili did not reveal significant differences that could explain their distinct stability. OutG shares over 77% sequence identity with PulG (**Figure 5A**), with three regions showing sequence divergence, listed here from the most to least divergent: the 12-residue C-terminal extension (CTer) present only in OutG; the N-terminal part of the α3-β1 loop comprising residues 69-83 (L) and the region corresponding to the calcium-binding site comprising residues 114-125 (Ca) (**Figure 5A, B**). To identify determinants of pilin stability, we designed different chimeras of OutG and PulG (**Figure S4**) and assessed their structural and thermal stability *in vitro* as well as their assembly into pili *in vivo*.

**Figure 5:**
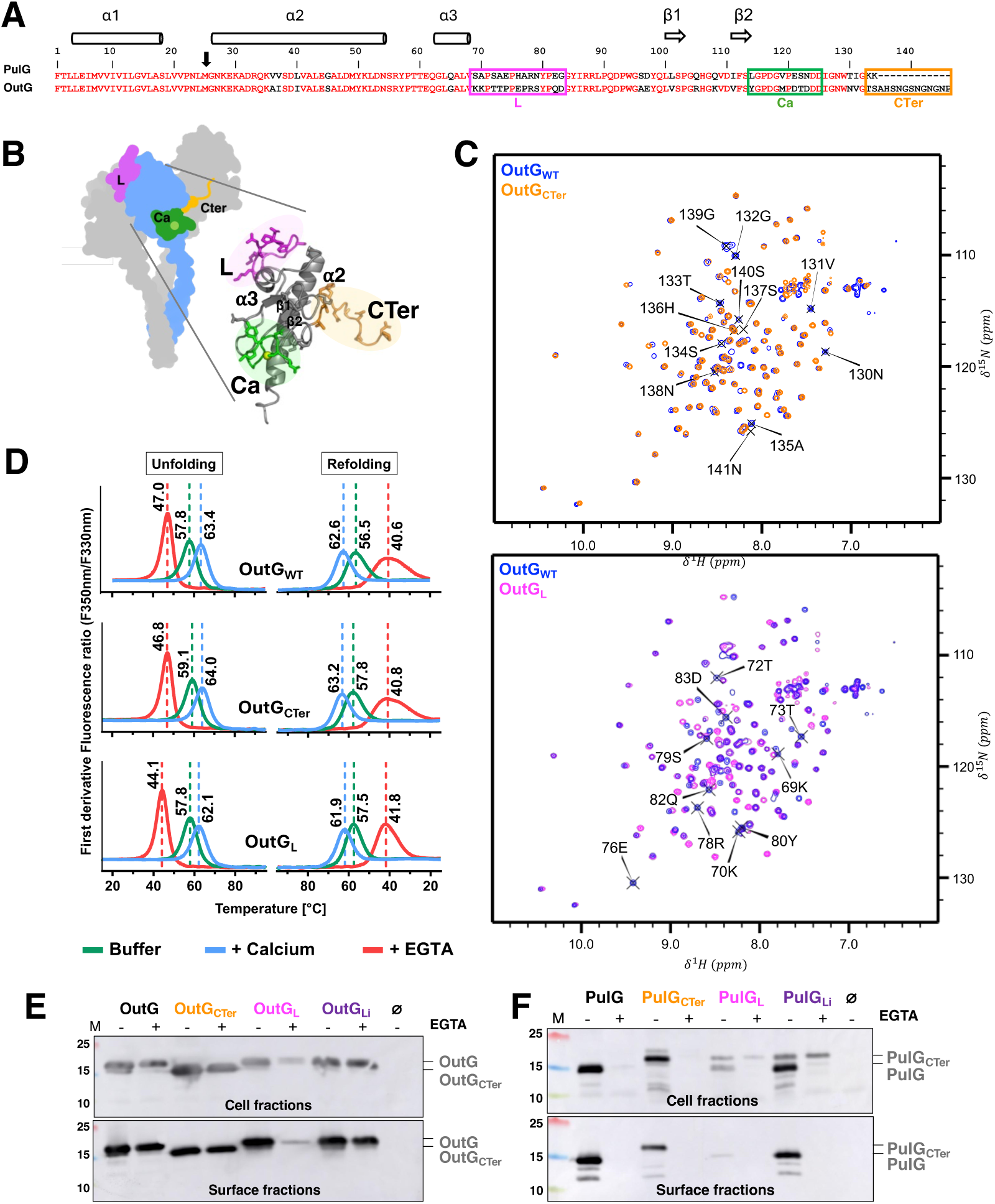
The α3-β1 loop (L) and the C-terminal extension (CTer) of OutG are not required for its stability. **A.** Sequence alignment of *K. oxytoca* PulG and *D. dadantii* OutG major pilins. The coloured frames indicate variable regions: L (loop, magenta), Ca (calcium binding site, green) and CTer (C-terminus, orange). Secondary structures are shown on top. *In vitro* studies were performed with soluble periplasmic domains starting from the black arrow. **B**. Structural organisation and surface exposure of OutG pilin variable regions in the monomer (bottom) and within the pilus (top), colour code as in A. **C.** Superimposed ^1^H-^15^N HSQC spectra of OutG and its variants. Resonance assignments of substituted residues are labelled. **D**. Temperature-induced unfolding and refolding patterns of OutG variants. First derivatives of 350 nm/330 nm fluorescence ratio with respect to temperature are presented. The transition temperature (Tm) values are indicated above each curve. All protein samples in C et D are at 0.075 mM in 50 mM HEPES,100 mM NaCl, pH 7 buffer (green curves), supplemented with 10 mM CaCl_2_ (blue curves) or 5 mM EGTA (red curves). **E.** Immunodetection of OutG, OutG_CTer_, OutG_L_ and OutG_Li_ variants in the cell- and surface fractions of bacteria harbouring the T2SS in the absence (-) or presence (+) of 2 mM EGTA. Ø indicates strain harbouring an empty vector. **F.** Immunodetection of PulG, PulG_CTer_, PulG_L_ and PulG_Li_ as in E.

We first focused on the C-terminal extremity as the most divergent region between OutG and PulG; OutG presents a 12-residue extension and lacks positively charged residues contrary to PulG, which ends with two lysine residues (**Figure 5A**). These regions are disordered and dynamic in both monomeric pilin and assembled pilus structures (**Figure 5B**). To assess their role in protein stability, we studied the variant OutG_CTer_ containing the two C-terminal residues of PulG, instead of its 12-residues C-terminus **(Figure S4)**. As shown in the ^1^H-^15^N HSQC NMR spectrum, the overall fold of the variant is preserved (**Figure 5C)**. Spectral superposition revealed only local perturbations around the mutated residues. Consistently, the Tm and the shape of thermal unfolding/refolding curves remained comparable to those of wild type OutG (**Figure 5C top, 5D**). Furthermore, calcium depletion did not affect the assembly of OutG_CTer_ variant into pili *in vivo* (**Figure 5E**), nor the stability of these pili *in vitro* (**Figure S5)**. Conversely, the PulG_CTer_ variant with the C-terminal extension of OutG was as unstable as PulG_WT_ and therefore did not assemble into pili in the presence of EGTA (**Figure 5F**). Together these results show that the C-terminal extremity (CTer) is not involved in pilin nor pilus stability.

The divergent α3-β1 loop L contains the same number of residues in both pilins, but it is more positively charged in OutG than in PulG (**Figure 5A, S4**). In the cellular context, the OutG_L_ variant containing the L sequence of PulG (residues 69-83, **Figure S4**) showed reduced stability and did not assemble into pili in the presence of EGTA **(Figure 5E)**. *In vitro*, the monomeric subunit of this variant showed a slight decrease of thermal stability in the presence of EGTA, with a Tm of OutG_L_ lower by ∼3°C compared to OutG_WT_ (**Figure 5D**). While more sensitive than the wild type, the fold and stability of this OutG_L_ variant were not as drastically dependent on calcium as PulG. This is consistent with the observation of only local NMR spectral perturbations around the mutated region in the OutG_L_ variant compared to wild type (**Figure 5C, bottom**).

*In vivo*, the PulG_L_ variant carrying the loop L from OutG (residues 69-83) was more unstable and barely detectable even in the presence of calcium (**Figure 5F**). Since the chimera of the L region contain residues 82 and 83 implicated in pilin interfaces, we also generated chimera designated Li wherein the swaps only included residues 69-81. Interestingly, these chimeras recapitulated the stability of their native counterparts, with OutG_Li_ being as stable as OutG_WT_, and PulG_Li_ being degraded like PulG_WT_ in the absence of calcium (**Figure 5E, 5F**). Therefore, we concluded that the divergent Li region comprising residues 69-81 is not involved in pilus stability.

The calcium-binding sites display a similar sequence in both proteins with small variation (**Figure 5A**), including two additional aspartate residues (D121 and D123) in OutG, though their side chains are not involved in the calcium coordination. We swapped the segments comprising residues 114 to 123, introducing five relatively conservative substitutions into OutG or PulG (**Figure S4**). Strikingly, these replacements affected the stability of both pilin subunits and assembled pili. In the presence of EGTA, the OutG_Ca_ variant containing the calcium-binding site of PulG was less stable than OutG_WT_ and did not assemble into pili (**Figure 6A**). Conversely, compared to PulG_WT_, substitution of the calcium-binding site in PulG_Ca_ improved its stability *in vivo* and enabled pilus assembly on the cell surface, regardless the presence of calcium ions (**Figure 6B**). *In vitro,* OutG_Ca_ showed a significantly lower thermal stability than OutG_WT_ and closer to that of PulG in all conditions, including in the presence of additional calcium. Importantly, OutG_Ca_ was unable to refold in the presence of EGTA, unlike OutG_WT_ (**Figure 6C**). These results show that the calcium binding site is the major determinant of stability of both pilin and pili. NMR spectral perturbations of OutG_Ca_ beyond the mutation site, affecting the upstream β-sheet, support a stabilizing structural role for the calcium binding site (**Figure 6D**, **6E**).

**Figure 6:**
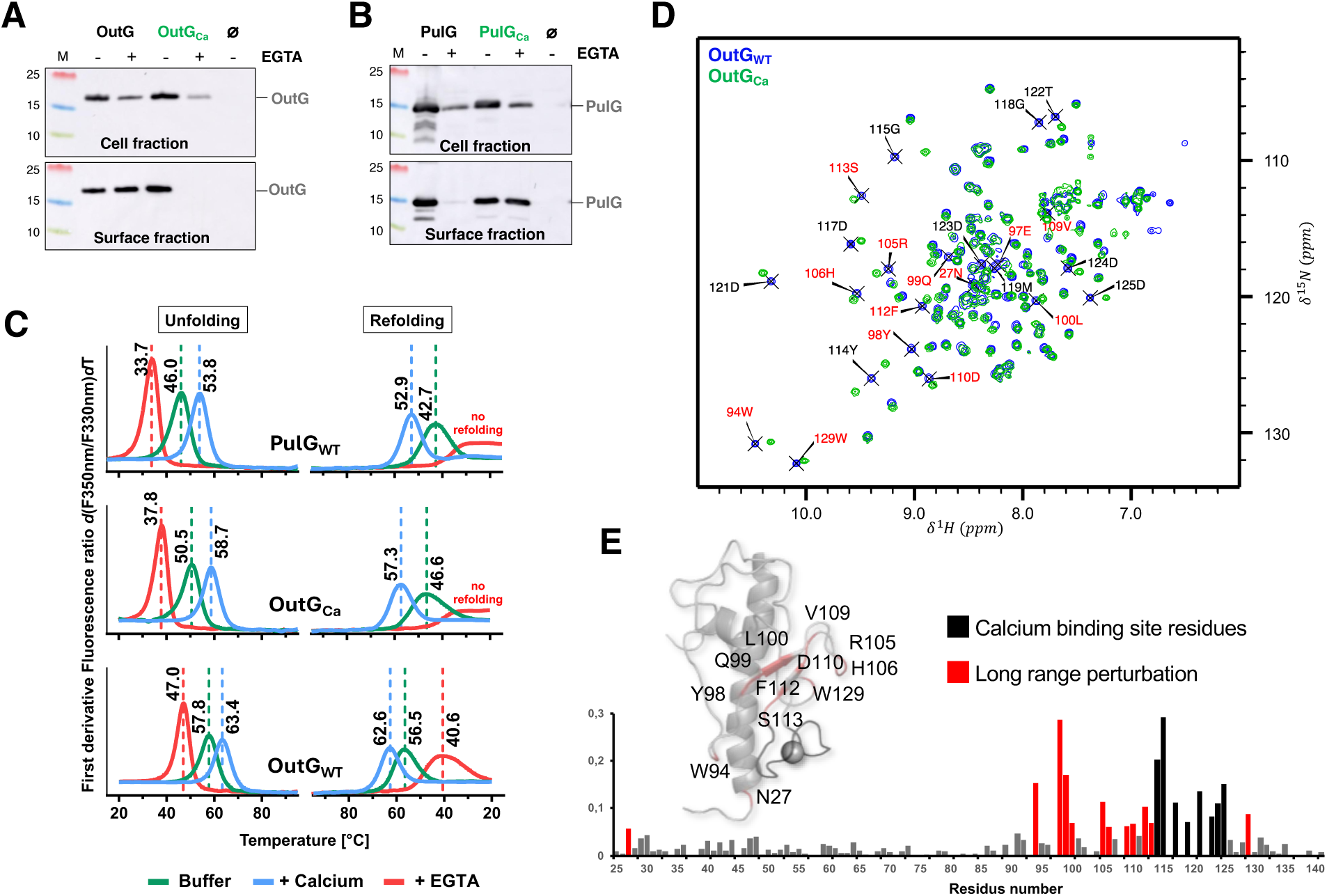
The calcium binding site determines pilin and pilus stability. Immunoblot analysis of pilus assembly from **A**. OutG_WT_ (OutG) and OutG_Ca_ variants and **B.** PulG_WT_ (PulG) and PulG_Ca_ variants in cell and surface fractions of bacteria cultured in the absence (-) or presence (+) of 2 mM EGTA. Ø indicates strain harbouring an empty vector. **C.** Temperature-induced unfolding and refolding patterns of PulG_WT_, OutG_WT_ and OutG_Ca_. The first derivative of 350nm/330nm fluorescence ratio with respect to temperature is shown. The transition temperature (Tm) values are indicated above each curve. **D.** ^1^H-^15^N HSQC of OutG_WT_ (blue) and OutG_Ca_ (green). The residues showing significant chemical shift changes are indicated. The calcium binding site residues are in black; residues with high CSP (> 0.05 ppm) and not belonging to the calcium binding site are in red. **E**. Chemical shift perturbation upon OutG_Ca_ mutation. Top: Ribbon presentation of OutG pilin structure coloured based on the CSP values.

### Secretion specificity determinants of T2SS endopilus

To evaluate the effect of the cellular context on pilin stability and function, we analysed the behaviour of pilin chimeras in the *D. dadantii ΔoutG* mutant. Upon growth on solid medium, all pilins generated comparable levels of T2SS pili (**Figure S6**). However, upon calcium depletion, the levels of PulG pili were strongly reduced, in contrast to those with OutG. Substituting the variable regions of PulG with their OutG counterparts confirmed the stabilizing effect of the calcium binding site. In the presence of EGTA, PulG_Ca_ pili were more abundant than those with PulG_WT_, while assembly of OutG_Ca_ pili was abolished (**Figure S6**). Notably, in *D. dadantii*, the pili produced by the chimeras carrying heterologous loop region (OutG_L_ and PulG_L_) showed stability comparable to that of their respective wild type pilins (**Figure S6**). This contrasts with the behaviour of these chimeras in the context of *K. oxytoca* T2SS reconstituted in *E. coli* (**Figure 5E, F**).

To analyse the ability of pilin variants to promote secretion, *D. dadantii ΔoutG* cells expressing pilin of interest were grown in liquid media. Culture supernatants (S) containing secreted proteins were separated from the cells (C) and both fractions were probed by Western blot with antibodies against one of the substrates of the Out T2SS, the pectate lyase PelD (**Figure 7**). In the presence of OutG_WT_, more than 50% of PelD was detected in the supernatant, while only trace amounts (<10%) of PelD were observed with PulG_WT_, comparable to cells harbouring an empty vector. This suggests that PulG lacks key determinants required for recognition of PelD or is improperly integrated in the Out system.

**Figure 7.**
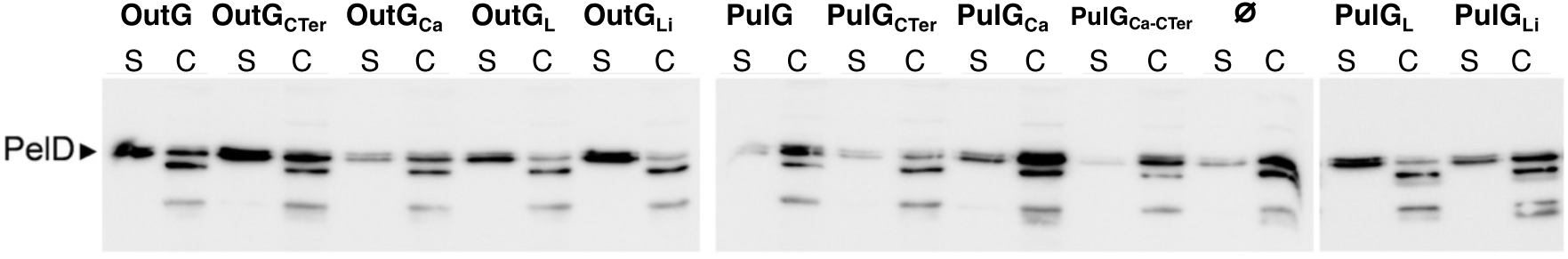
Secretion assay with wild type PulG and OutG and their chimeras expressed in *D. dadantii ΔoutG*. Bacterial culture supernatants (S) were separated from the cell fractions (C) and probed by Western blot with anti-PelD. Ratio of PelD in “S” *versus* “C” fraction reflects the ability of each pilin variant to promote secretion. Ø indicates strain harbouring an empty vector.

Addition of the long C-terminal extension of OutG into PulG did not improve PelD secretion with the resulting PulG_CTer_ variant, and conversely, the reciprocal substitution in OutG (OutG_CTer_) did not impair its secretion. Similarly, the calcium-binding site substitution in PulG_Ca_ did not increase the PelD ratio in supernatant yet it increased the total amount of produced PelD (**Figure 7**). Combination of two substitutions, the calcium-binding site and the C-terminal extension in PulG_Ca-CTer_ lowered the PelD level but did not improve its secretion, indicating that these two regions do not confer functional specificity in protein secretion.

In contrast, substitution of the α3-β1 loop (residues 69-83) improved PelD secretion with PulG_L_. Consistently, the amount of secreted PelD was reduced with OutG_L_ variant. Notably, PulG_Li_ and OutG_Li_ variants showed milder phenotypes, with lower secretion efficiency with PulG_Li_ and a wild-type level of secreted PelD with OutG_Li_ (**Figure 7**). Together these data indicate that the variable L region plays a role in specific secretion in *D. dadantii*.

## Discussion

Despite sharing high sequence identity, conserved fold and helical assembly mode, OutG and PulG exhibit significant differences in pilin and pilus stability. In the absence of calcium ions, PulG monomer adopts multiple disordered conformations, is completely degraded *in vivo*, and its pilus is dissociated *in vitro*^14^. OutG pilin and pilus, however, remain stable regardless of calcium availability (**Figure 1B, 2A**). We showed that the calcium-binding site is critical for the stability of both pilins and assembled pili. Its exchange between OutG and PulG was sufficient to alter protein stability and pilus assembly (**Figure 6, S6**), despite conserved overall structure. Furthermore, high CSPs detected beyond the mutated calcium-binding site **(Figure 6E),** reflect the loss of some stabilizing intramolecular contacts, such as Q99-Y114, and a more general structural role of the calcium binding site. Its variation between species suggests an evolutionary adaptation to specific environment. Indeed, the sequences of the calcium binding site vary across species, forming even in some cases additional structural elements such as a short α helix in *Vibrio cholerae* EpsG^13^ (**Figure S7**).

Importantly, despite similar pilin structures and helical assemblies, the surface-exposed polarities of PulG and OutG pili differ significantly. PulG endopili display a strongly negatively charged surface favouring calcium attraction, whereas OutG endopili exhibit a more balanced electrostatic distribution with localized positively charged patches **(Figure 4C)**. These differences could be related to their distinct substrate specificities. Functionally, the PulG endopilus is specialized for secreting a single substrate, the pullulanase^15^, while the OutG endopilus mediates the secretion of large amounts of multiple plant cell wall-degrading enzymes, mainly polysaccharide lyases^22^ with diverse polarities and as-yet-unidentified recognition signals. These enzymes belong to ancestrally and structurally diverse polysaccharide lyase families (PL1, PL3, PL9 and PL10, www.cazy.org). However, all these enzymes possess a calcium in their catalytic sites^23–26^ and the structure of some of them is stabilized by additional calcium ions^25,27^. Therefore, they may compete with OutG for calcium during their folding and transit through the *D. dadantii* periplasm. Given that T2SS models propose a direct interaction between the endopilus and its substrate during secretion^1^, OutG may have evolved reduced calcium dependence and higher overall stability to resist fluctuating periplasmic calcium levels. Interestingly, periplasmic pectate lyases of the PL2 family, which are not secreted and permanently reside in the periplasm to degrade pectic oligomers entering the cells, contain in their catalytic site a transition metal either Co^2+^, Ni^2+^ or Mn^2+^ instead of calcium^28,29^. This adaptation may avoid competition for calcium in the periplasm, illustrating an evolutionary strategy to optimize metal availability. In contrast, during *Klebsiella* infection, calcium would not be limiting as it is available at millimolar concentrations in airway mucus, tissues and blood^30^. This lack of selective pressure for calcium availability may have led PulG to become strongly calcium dependent. Furthermore, this dependence on calcium may represent an added regulatory layer of T2SS assembly and function *in vivo*^31^.

Secretion assays in *D. dadantii* demonstrate that major pilins also encode determinants of substrate or secretion specificity, challenging the conclusions of some previous studies^32^. Our work shows that PulG does not promote protein secretion in the *D. dadantii ΔoutG* mutant under physiological conditions and expression levels. In addition, the reduced PelD levels observed with OutG_L_ and OutG_Ca_ variants (**Figure 7**, S + C fractions) suggests that heterologous pilins could improperly interact with T2SS components and/or substrates causing partial degradation of the latter.

Swapping the divergent L loop revealed unexpected *in vivo* behaviours, differentially affecting pilin stability depending on the host T2SS system and the nature of residues 82 and 83. Although this loop is surface exposed, residues Q82 and D83 are buried in the OutG pilus, with D83 forming a salt bridge with K108 of the P+1 subunit (**Figure 3G**), an interaction absent in PulG. Consistently, PulG_L_ uniquely restores PelD secretion in *D. dadantii* (**Figure 7**), with full complementation by PulG_L_ and partial by PulG_Li_. This functional role may result from the interactions of this variable loop with other T2SS components or substrates. Further analyses are needed to identify these partners and elucidate how this loop modulates substrate specificity.

Overall, our results demonstrate that two variable regions of major pilins, the calcium-binding site and the α3-β1 loop, fulfil distinct functional roles: the calcium-binding site contributes to pilin and pilus stability, whereas the α3-β1 loop is critical for secretion specificity. Although T2SSs are relatively promiscuous, and can indeed assemble heterologous pilins, their sequence determinants are fine-tuned during evolution to ensure optimal secretion under physiological conditions in their specific environment. Surprisingly, we found that the long C-terminal extension, a distinctive feature of OutG, does not affect pilus stability or PelD secretion. This extension might instead play a role in capture and transport of other substrates among ∼20 proteins secreted by the Out T2SS in *D. dadantii*.

## Materials and Methods

### Bacterial strains, plasmids and molecular biology techniques

The list of *Dickeya dadantii* and *Escherichia coli* strains and plasmids is provided in **Table S4**. *D. dadantii* non-polar *ΔoutG* mutant was constructed by marker exchange-eviction mutagenesis^33^. An *nptI-sacB-sacR* cassette was introduced into unique *Eco*RI site of the *outH* gene onto pGM-T*outG-H-I* plasmid and next, introduced into the *D. dadantii* A5652 chromosome by homologous recombination. The resulting A3649 strain was Km^R^, sucrose-sensitive and secretion deficient. The *outG* allele carrying TAG and TGA Stop codons in place CAG and CGA codons encoding Gln5 and Arg6 was then exchanged for the chromosomal allele of A3649 by selection for sucrose tolerance. The correctness of recombination of *outG* mutant allele in the resulting A6408 strain was checked by sequencing.

*E. coli* strain DH5α was used for cloning. Assays of pilus assembly *via* the *K. oxytoca* T2SS were performed in *E. coli* PAP7460 carrying the complete *K. oxytoca pul* gene cluster with the *pulG* deletion on plasmid pCHAP8184 and complemented with different *pulG* or *outG* alleles on plasmid pSU18. Pili were purified from strain PAP7501.

*E. coli* and *D. dadantii* strains were cultured in LB medium supplemented with antibiotics as required: carbenicillin 100 μg.mL^-1^; chloramphenicol 25 μg.mL^-1^; kanamycin 30μg.mL^-1^. The details of plasmid constructions are provided in the Supplementary Methods. The list of oligonucleotides is provided in **Table S5**.

### Expression and purification of soluble domains of major pilins

Sequence of the mature soluble domain of OutG (residues M25-P146) (**Figure S4**) was cloned into pET20b^+^ vector with an N-terminal fusion comprising the pelB signal peptide for periplasmic targeting, followed by a hexa-histidine tag and a TEV protease cleavage site. From this expression plasmid of wild-type OutG were constructed the mutant plasmids encoding OutG_Ca_ (Y_114_-D_123_ region replaced by L_114_-N_123_ region comprising the calcium binding site of PulG) and OutG_L_ (K_69_-D_83_ region replaced by S_69_-G_83_ region forming the loop L of PulG) (**Table S3**).

Unlabelled proteins were produced in LB medium, and uniformly ^15^N and ^15^N/^13^C labelled proteins were produced in M9 minimal medium using 1 g/L of ^15^NH_4_Cl and 4 g/L ^13^C glucose, as the sole nitrogen and carbon sources, respectively. Protein expression of OutG was induced by addition of 1 mM IPTG for 3 hours at 37°C in *E. coli* BL21 Star (DE3) pLysS strain. Bacteria were harvested by centrifugation at 8,000 × g for 10 min, resuspended in lysis buffer (50 mM Tris-HCl pH 8.0, 100 mM NaCl) supplemented with antiprotease inhibitor cocktail (cOmplete™ EDTA-free, Roche). Bacteria were lysed by sonication, lysate was then clarified by centrifugation at 16,000 × g for 1 hour, and supernatant was finally filtered through 0.22 µm membrane. Proteins were purified by immobilized metal affinity chromatography onto an HiTrap Chelating HP column (Cytiva) equilibrated in 50 mM Tris-HCl pH 8.0, 100 mM NaCl, 10 mM imidazole. Bound proteins were eluted using a linear gradient to 500 mM imidazole and subsequently incubated overnight with TEV protease to remove the His-tag. The mixture was passed over a second HiTrap column, and the flow-through containing cleaved OutG was concentrated using 5 kDa cut-off Vivaspin devices (Cytiva). Final size exclusion chromatography was performed with a HiLoad 16/600 Superdex 75 pg column (Cytiva) equilibrated in 50 mM HEPES pH 7.0, 100 mM NaCl. All purification steps were performed at 4°C and evaluated by SDS-PAGE analysis. All purification buffers were supplemented with EDTA-free protease inhibitor cocktail (cOmplete™ EDTA-free, Roche).

PulG expression and purification were performed as previously described but with lysis performed through sonication instead of osmotic shock^34,14^.

### SDS-PAGE and Western blot analysis of bacterial extracts

Total bacterial extracts and sheared pilus fractions were analysed by denaturing sodium dodecyl sulphate (SDS) polyacrylamide gel electrophoresis (PAGE) on 10% Tris-Tricine gels^35^. Proteins were transferred by semi-dry method in the 1-step rapid transfer buffer (Thermo) using the Power blotter (Thermo) onto nitrocellulose membranes (0.2 μm pore size). The membranes were blocked for 1 hour at room temperature (RT) in 5% low-fat dry milk in Tris-buffer saline buffer with 0.05% Tween-20 (TBST). The membranes were incubated for 1 hour with the primary polyclonal PulG or OutG antibodies pre-adsorbed with the extract of PAP7460 (pCHAP8184) strain and diluted to 1:2000 in TBST, followed by three 10-min washes in TBST. Upon 1 hour of incubation in TBST with the secondary goat anti-rabbit antibodies coupled with horse-radish peroxidase (HRP) at 1:10 000 dilution, and four 10-min washes in TBST, the membranes were developed with ECL or ECL-2 (Thermo). Chemiluminescent and fluorescent signals were recorded on Amersham AI680 and Typhoon SLA-9000 (GE) imagers, respectively. The images were visualized with Affinity Photo and quantified with ImageJ.

### Preparation of pili

The OutG and PulG pili were purified from strain PAP7501 containing plasmid pCHAP8184 and pMS1100 (OutG_WT_), pMS1413 (OutG_CTer_) or pCHAP8658 (PulG_WT_). Bacteria were cultured for 5 days on 30 LB plates supplemented with Amp, Cm and 0.2% D-maltose. Bacteria were collected and gently resuspended in LB. To shear the pili from the cell surface, bacterial suspensions were vortexed for 2 min and passed 6 times through a 23-Gauge needle. Bacteria were pelleted for 15 min at 9,000 rpm in 50-mL Falcon tube rotor in the Eppendorf centrifuge 5810R at 4°C. The supernatants were further subjected to centrifugation for 15 min and 15,000 × g. Cell-free supernatants were subjected to ultracentrifugation for 1 hour at 45,000 rpm (81,648 × g) in a Type 60 Ti rotor (Beckman). The pellets containing the crude surface fractions were resuspended in 200 μL of cold 50 mM HEPES, pH 7.2, 50 mM NaCl, pH 7.2 and stored at 4°C until further use.

### Pilus assembly and stability assays

For pilus assembly assays in *E. coli* bacteria were cultured for 48 hours at 30°C on LB agar containing appropriate antibiotics and 0.2% D-maltose, supplemented or not with 2 mM EGTA. Bacteria were suspended in LB, normalized to OD_600nm_ of 1 and vortexed for 1 min to shear the surface pili. After a 10 min centrifugation at 16,000 x g at 4°C, the supernatants were removed, and bacterial pellets were suspended in 100 μL of SDS sample buffer. The supernatants were further cleared by a 10-min centrifugation 16,000 x g and 0.7 mL was removed off top and transferred to a new 1.5 mL eppendorf tube. Proteins were precipitated in 10% tri-chloro-acetic acid for 30 min on ice followed by a 30-min centrifugation at 21,000 x g. The pellets were washed twice in cold acetone, air-dried and resuspended in 70 μL of SDS sample buffer. Equivalent volumes (typically 3 μL) of each fraction were analysed by denaturing electrophoresis on Tris-tricine SDS-PAGE, followed by western blot.

To test the stability of purified pili, 20 μL of each pilus preparation was mixed with 5 μl of either water, 25 mM EGTA or 25 mM CaCl_2_ and incubated under static conditions in the water bath at 60°C for 1 hour. After a 30-min ultracentrifugation at 50, 000 rpm in a TLA-55 rotor, supernatant fraction was removed and mixed with an equal volume of 2X SDS sample buffer. The pellets were resuspended in 25 μL of SDS sample buffer. The equivalent amounts of pilus and supernatant fractions were analysed on SDS-PAGE and stained with Coomassie blue R. The intensity of the bands was quantified using ImageJ. GraphPad Prism was used for graph plotting and statistical analysis using nonparametric ANOVA test with multiple comparisons.

### Thermal stability analysis by NanoDSF

Thermal stability of proteins was assessed using a Prometheus NT.48 instrument (*NanoTemper Technologies GmbH*). All protein samples were prepared at a final concentration of 75 µM in their respective SEC buffer in three conditions: buffer alone, buffer supplemented with 10 mM CaCl₂, or buffer supplemented with 5 mM EGTA. Triplicates of 10 µL of each sample were loaded into standard capillaries and heated from 15-20°C to 90-95°C, at a rate of 1°C·min⁻¹. Excitation power was set to 20% to maintain fluorescence intensities below 10,000 counts. Intrinsic fluorescence at 330 nm and 350 nm was recorded simultaneously, and unfolding transitions were monitored using the F_350_/F_330_ fluorescence ratio. Both heating and cooling phases were recorded to evaluate reversible unfolding. Data analysis and melting temperature (Tm) calculation were performed using PR.thermControl software (NanoTemper Technologies).

### NMR experiments

NMR spectra for resonance assignments were acquired with ^15^N/^13^C labelled samples between 0.075 and 0.5 mM in 50 mM HEPES, pH 7.0, 100 mM NaCl at 25°C on a 600 MHz Avance III HD and a 800 MHz Avance NEO spectrometers (Bruker Biospin, Billerica, USA) both equipped with a cryogenically cooled triple resonance ^1^H [^13^C /^15^N] probe (Bruker Biospin, Billerica, USA). The pulse sequences were employed as implemented in the TOPSPIN 4.2 (Bruker, Biospin, Billerica, USA) and IBS libraries^36^. OutG wild type ^1^H, ^15^N, and ^13^C backbone and side-chain resonances were assigned in our previous work and were deposited to the BMRB under accession number 51296^19^. OutG calcium-free state resonances were assigned by using standard set of experiments as previously described^19^. Calcium-free NMR samples were prepared by dialysing purified proteins against buffer supplemented with 10 mM EGTA. 2,2-Dimethyl-2-silapentane-5-sulfonate (DSS) signal was taken as 0 ppm for referencing proton chemical shifts and ^15^N and ^13^C chemical shifts were indirectly referenced to DSS^37^. CcpNmr Analysis^38^ was used for NMR data analysis.

### NMR chemical shift perturbation analysis

OutG calcium binding and mutations effects were monitored by comparison of the ^1^H-^15^N HSQC spectra at 25°C. Chemical shift perturbations (CSP) of backbone amide resonances were quantified by using the equation CSP = [ΔδH^2^ + (ΔδN∗0.159)^2^]½, where ΔδH and ΔδN are the observed ^1^H and ^15^N chemical shift changes between the two experimental conditions. Residues displaying statistically significant CSP values (higher than the average plus one standard deviation) were considered for the analysis as described in^39^.

### NMR structure calculation

The structure of the periplasmic domain of OutG was determined by iterative structure calculation with ARIAweb/CNS^17,40^ making use of the standard torsion angle simulating annealing protocol. An AlphaFold3^41^ model was used as initial structure to guide the assignment of the 3D ^15^N-NOESY and ^13^C-NOESY cross peaks, only for the first iteration. Restraint combination^42,43^ was used for the first 4 iterations. A calcium ion (Ca^2+^) was included and coordinated with distance restraints involving residues identified from the Ca^2+^/EGTA CSP data (side chain oxygen D117 and D125; backbone carbonyl oxygen of residues Tyr114 and Met119). Proline chemical shift analysis with Promega^44^ had revealed that proline residues at positions 75 and 103 were predicted to be in cis conformation which was imposed in the structure calculation. Backbone dihedral angles were predicted based on assigned chemical shifts of OutG by using TALOS-N^18^. The dihedral angles categorised as “Strong” or “Generous” were included as restraints. Nine iterations were used in the structure calculation with calculation of 50 conformers, and the 10 lowest energy ones used for analysis. After the final iteration, the lowest energy conformers were refined in an explicit water shell^45^. Structure quality was evaluated with the PSVS server^46^.

### Cryo-EM data collection and image processing

Lacey carbon film 300 mesh cooper grids (AGS166-3, Agar scientific) were glow-discharged (15 mA for 25 sec) immediately prior to pili sample application. Aliquots of pili sample were applied to the grids, which were blotted with a blot force of 0 for 3 s at 4°C and 100% humidity, and flash-frozen in liquid ethane using a Vitrobot Mark IV (Thermo Fisher Scientific). Cryo-EM data were collected on a Titan Krios G3 TEM (Thermo Fisher Scientific) equipped with a Falcon 4i direct electron detector (Thermo Fisher Scientific) and a Selectris X energy filter. Images were recorded through EPU software in electron counting mode at a nominal magnification of 165,000× (pixel size of 0.77 Å/pixel), with a total accumulated dose of 40 e⁻/Å², and with a defocus range from −0.8 to −2.4 µm.

Data processing was performed in cryoSPARC^47^. Frame shifts and beam-induced motions in the movies were corrected using the Patch Motion Correction tool, and the contrast transfer function (CTF) was estimated using the Patch CTF Estimation tool in cryoSPARC. Pili filaments were manually picked using the Manual Picker to generate filament templates, which were then used for automated picking with the Filament Tracer tool. The selected filaments were segmented into overlapping boxes and subjected to 2D classification to remove poorly aligned particles and false picks. The selected segments were used to calculate the average power spectrum, which was applied to determine the initial helical symmetry parameters. Using these parameters, helical reconstruction was carried out with the Helix Refine tool in cryoSPARC. Helical parameter refinement was performed at each iteration once the map resolution reached 6 Å. The resulting map was further improved through iterative rounds of per-particle CTF refinement and Helix Refine until no further resolution improvement was observed. Data processing statistics are provided in **Table S3.**

### Model building and refinement

AlphaFold^41^-predicted structures of OutG and PulG were first rigid-body docked into the respective cryo-EM maps and subsequently refined in ISOLDE^48^. The resulting models were manually adjusted in COOT^49^, followed by real-space refinement in Phenix^50^ to improve the overall map fitting and model geometry. Model building statistics are provided in **Table S3**.

### Pectinase secretion assay

A series of cultures of *D. dadantii* A6408 carrying pGM-T plasmids with various *outG* and *pulG* variants were grown in 4 mL of LB supplemented with 100 µg.mL^-1^ of ampicillin at 30°C for 8 hours to attain similar OD_600nm_ values. Then, 50 µL of these cultures were used to start a new series of cultures in 4 mL of LB supplemented with 0.2% galacturonate and 100 µg.mL^-1^ ampicillin at 30°C for 17 hours. Culture supernatants were separated from cells at 10,000 x g for 2 min and both fractions were loaded onto 10% SDS-PAGE and probed with antibodies against PelD at 1:1,500^27^ followed by secondary anti-rabbit goat IgG conjugated to peroxidase (Sigma-Aldrich, St. Louis, USA) at 1:20,000. Chemiluminescence signals were collected with Immobilon substrate (Millipore, Burlington, USA) using Fusion FX imaging system (Vilber Lourmat, Marne-la-Vallee, France). The data were collected from at least two technical replicates of three biological experiments and analysed with a two-sample Mann–Whitney test by comparing the values of supernatant (S) to cell (C) fractions. Graphs were generated using PRISM software. The ratio of PelD in S to C fraction reflects efficiency of secretion.

### Analysis of *D. dadantii* surface exposed pili

*D. dadantii* A6408 carrying pGM-T plasmids with various *outG* and *pulG* variants were grown in 4 mL of LB supplemented with 100 µg/mL ampicillin at 30°C for 8 hours to attain similar OD_600nm_ values. Then, 50 µL of these cultures were spread onto solid LB plates supplemented with 0.2% galacturonate, 1mM IPTG and 100 µg.mL^-^1 ampicillin and incubated at 30°C for 24 hours. Cells from ∼4 cm^2^ were collected, resuspended in 0.5 mL of LB, vortexed for 10 sec and pelleted at 5,000 x g for 3 min. The supernatant containing essentially the T2SS pili and flagella were separated by 15% SDS-PAGE and probed with a mixture of antibodies directed against OutG (1:4,000) (this study), PulG (1:3,000)^11^, and OmpA (1:10,000)^51^ adsorbed with cell lysate of A6408 strain. OmpA was used as a loading control. Chemiluminescence signals were collected and analysed as above with PelD.

### Structure visualisation and analysis

Structures were visualised and analysed with ChimeraX^52^ and PyMOL (Schrödinger, LLC). Surface electrostatics potential analysis was performed using the Adaptive Poisson-Boltzmann Solve, APBS^53^.

## Data availability

The solution structure of monomeric OutG is deposited in the Protein Data Bank under accession code 29JD. The cryo-EM maps of OutG and PulG pili have been deposited to the EMDB (codes EMD-75832 and EMD-75833, respectively) and the models are available at the PDB (accession codes 11ME and 11MF, respectively).

## Acknowledgments

This work was supported by Agence Nationale de la Recherche (grants Synergy-T2SS ANR-19-CE11-0020 and SecSPARC ANR-25-CE11-2325). RD was funded by Synergy-T2SS ANR-19-CE11-0020. SI was supported by the PAUSE Program (PAUSE-Solidarity with Ukraine) from Collège de France. TJ was funded by European Union’s Horizon 2020 Research and Innovation program under the Marie Sklodowska-Curie Grant Agreement No. 765042. EHE was supported by NIH GM122510. We thank Institut Pasteur technological platforms of: Production and Purification of recombinant Proteins for TEV production; Molecular Biophysics for access to the machines and particularly Sébastien Brûlé for his technical help; Biological NMR Technological Platform for the access to the NMR spectrometers as well as NanoImaging Core facility, created and supported by a PIA grant (EquipEx CACSICE: ANR-11-EQPX-0008). The 800-MHz NMR spectrometer of the Institut Pasteur was partially funded by the Région Ile de France (SESAME 2014 NMRCHR grant no 4014526). The authors acknowledge Youssef Ghorbal and Tru Quang Huynh for IT assistance.

## Author contributions

ML, SI, EHE, OF, VES, NIP conceived and designed the experiments. ML, SI, RD, BB, RRS, MV, TJ, OF, VES, NIP performed the experiments. ML, SI, RD, BB, RRS, TJ, EHE, MN, OF, VES, NIP analysed the data. ML, SI, RD, BB, RRS, EHE, MN, OF, VES, NIP wrote the manuscript.

## Declaration of Interests

The authors declare that they have no conflict of interest.

## Inclusion and diversity

We support inclusive, diverse, and equitable conduct of research.

## Notes

### Competing Interest Statement

The authors have declared no competing interest.

